# The burden of rare protein-truncating genetic variants on human lifespan

**DOI:** 10.1101/2020.06.02.129908

**Authors:** Jimmy Z. Liu, Chia-Yen Chen, Ellen A. Tsai, Christopher D. Whelan, David Sexton, Sally John, Heiko Runz

## Abstract

Genetic predisposition is believed to contribute substantially to the age at which we die. Genome-wide association studies (GWAS) have implicated more than 20 genetic loci to phenotypes related to human lifespan^1^. However, little is known about how lifespan is impacted by gene loss-of-function. Through whole-exome sequencing of 238,239 UK Biobank participants, we assessed the relevance of protein-truncating variant (PTV) gene burden on individual and parental survival. We identified exome-wide (P<2.5e-6) significant associations between *BRCA2, BRCA1, TET2, PPM1D, LDLR, EML2* and *DEDD2* PTV-burden with human lifespan. Gene and gene-set PTV-burden phenome-wide association studies (PheWAS) further highlighted the roles of these genes in cancer and cardiovascular disease as relevant for overall survival. The overlap between PTV-burden and prior GWAS results was modest, underscoring the value of sequencing in well-powered cohorts to complement GWAS for identifying loci associated with complex traits and disease.

## Main

Human lifespan is a heritable quantitative trait that reflects a range of health-related outcomes, environmental exposures, and chance. Twin and pedigree studies suggest that narrow-sense heritability of human lifespan ranges from 15-30%^2^, though these estimates may be inflated due to assortative mating^3^. The ability to identify genetic loci associated with lifespan has been limited by the lack of mortality information in most cohorts. Instead, to approximate lifespan, genome-wide association studies (GWAS) have used extreme longevity (e.g., those aged >90 years versus controls) or parental lifespan as phenotypes of interest. This has led to the identification of over 20 loci, several of which overlap age-related complex disease loci for Alzheimer’s disease (*APOE*), lung cancer (*CHRNA3/5*), cardiometabolic (*LPA, LDLR*) and immune-related disorders (*HLA, MAGI3*)^4–9^.

Rare protein truncating variants (PTVs) are reported to have an outsized effect on complex traits compared to common non-coding variants^10^ and are often poorly captured on GWAS genotyping arrays. PTVs typically shorten a protein’s coding sequence by introducing premature stop codons, frameshifts or aberrant splicing that lead to partial or complete loss of its function. Individuals who carry PTVs can be considered as “experiments of nature” that provide insights into gene function and that allow to extrapolate on efficacy and safety upon pharmacological inhibition of a gene product by drugs^11,12^.

Using whole-exome sequencing (WES) data from 302,331 UK Biobank participants, we sought to assess the impact of gene PTV burden on individual and parental survival using Cox proportion hazards models. We chose survival rather than age-at-death directly since this allowed us to account for censored data and most participants in the UK Biobank are still alive. After quality control (see Online **Error! Reference source not found.**), 238,239 individuals were available for analysis, of which 9,405 were deceased with their age at death recorded at the censoring dates. Fathers’ and mothers’ age at death were reported for 178,443 and 145,281 participants respectively, while 56,706 and 89,686 participants reported the age of their fathers and mothers respectively as being alive at the time of recruitment or follow-ups.

WES data were generated at the Regeneron Genetics Center as part of a precompetitive consortium effort^13^. We annotated PTVs using Variant Effect Predictor v96^14^ and the LOFTEE plugin^12^ and identified 572,780 high-confidence predicted PTVs with minor allele frequency <1%, including 386,785 singletons, in the canonical transcripts of 19,094 genes. Each individual carried on average 129 PTVs, which is consistent with previous estimates^12,15,16^. For each gene, we collapsed the PTVs into a single score, and performed burden survival analysis by testing the association between this score against six survival outcomes: individual survival, individual survival in males, individual survival in females, mother’s survival, father’s survival, and mother plus father combined survival (see Online Online Methods).

We identified seven genes associated with at least one survival outcome at exome-wide significance (P<2.5e-6) (Figure 1, Table 1). The strongest signal was observed for *BRCA2*, where PTV-burden was associated at exome-wide significance with reduced lifespan in five of the six outcomes analyzed (Figure 2): across all individuals (hazard ratio, HR=2.57, P=5.02e-15), in both females (HR=3.12, P=1.68e-10) and males (HR=2.25, P=8.43e-7) separately, and for combined parental (HR=1.60, P=2.30e-22) as well as mothers’ lifespan (HR=1.60, P=4.70e-26). *BRCA2* PTV burden was nominally associated with fathers’ lifespan (HR=1.21, P=1.35e-5). Similarly, PTV-burden in *PPM1D* reduced lifespan across all individuals (HR=3.14, P=7.70e-9) as well as in both females (HR=4.10, P=8.48e-6) and males (HR=2.73, P=8.07e-5) (Table 1, Supplementary Figure 1). In contrast, *BRCA1* and *TET2* PTV-burden associations with survival were sex specific, with *BRCA1* associated with female (HR=4.38, P=1.01e-6) and mothers’ survival (HR=1.64, P=5.35e-II), while *TET2* was associated with male survival (HR=1.92, P=6.44e-7). PTV-burden in *LDLR* was associated with survival in the combined parental analysis (HR=2.28, P=5.14e-7), and nominally significant in both mothers’ (HR=1.62, P=0.00197) and fathers’ survival (HR=1.60, P=0.0021). The *EML2* association signal was driven by males (HR=3.35, P=3.33e-8 versus P=0.21 in females), while *DEDD2* was significant for survival in mothers (HR=4.20, P=2.28e-7 versus P=0.55 in fathers).

**Figure 1.**
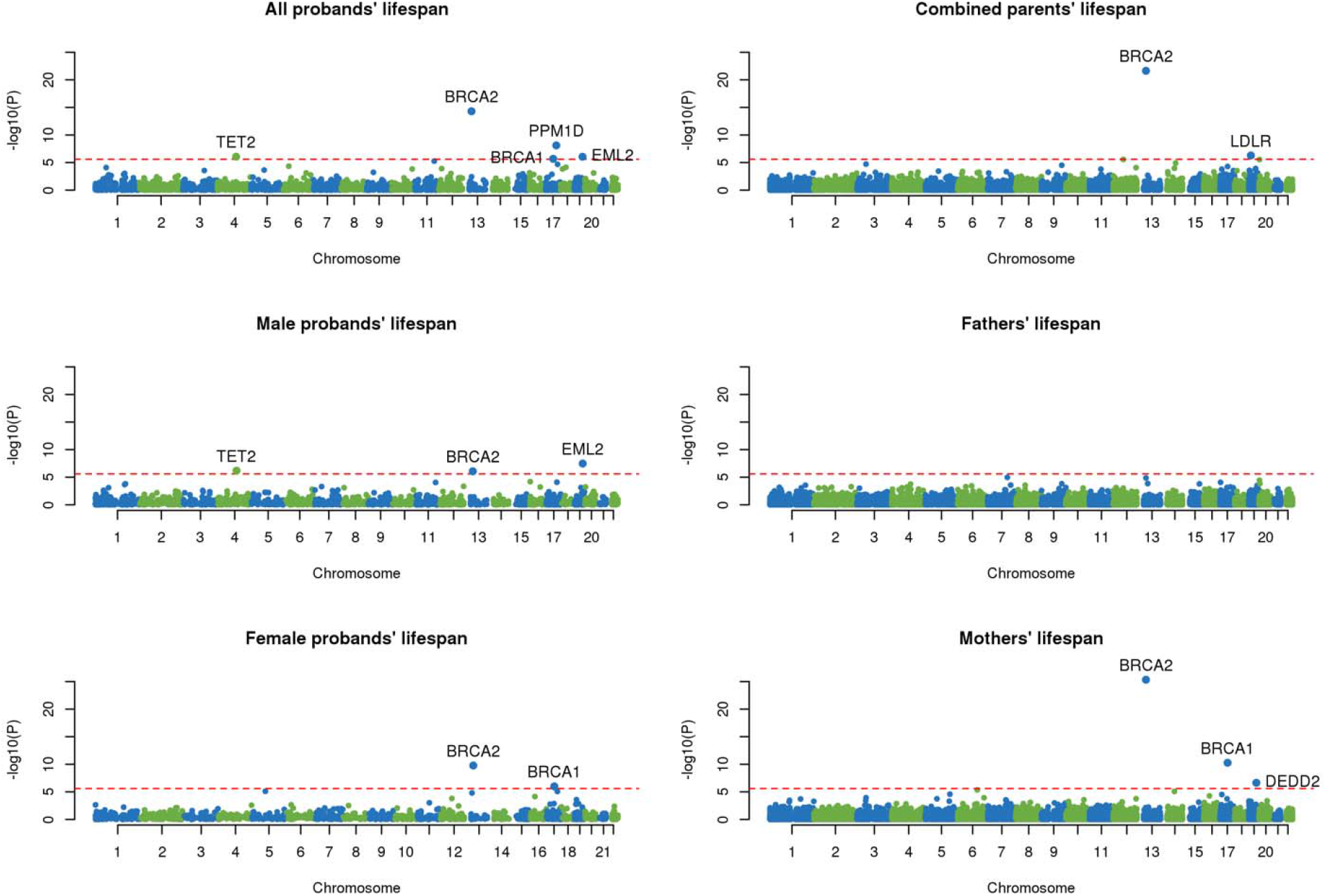
Manhattan plots of gene-based PTV-burden survival analyses in 238,239 UK Biobank participants and parents for six survival phenotypes analyzed. Each point represents a gene. The red-dashed line indicates the exome-wide significant threshold of P < 2.5e-6. Genes exceeding this threshold are labeled.

**Figure 2.**
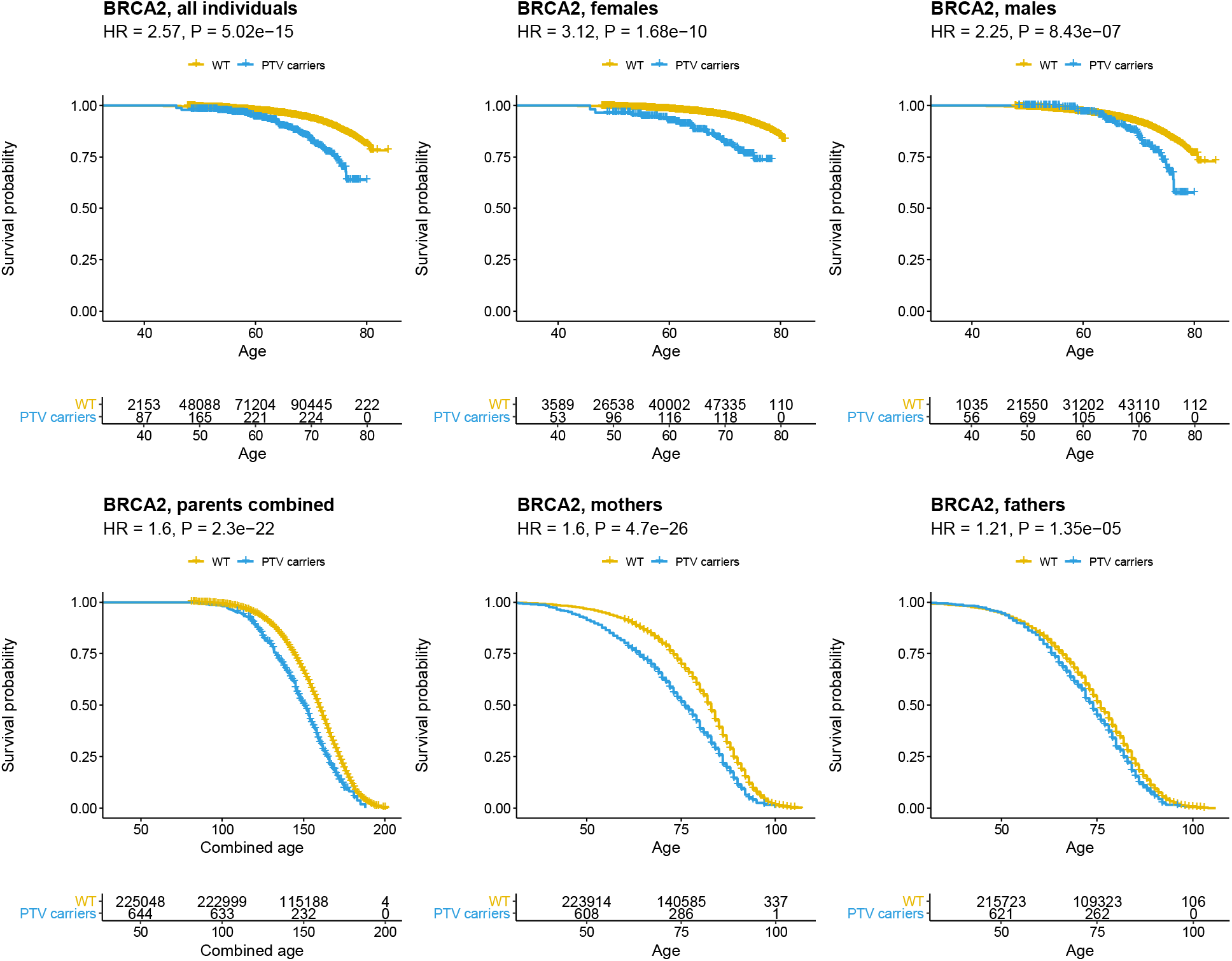
Kaplan-Meier curves for BRCA2 PTV-burden survival analyses across the six survival phenotypes analyzed in this study. Each cross represents a right censored observation. The shaded areas represent the 95% confidence interval of the curve. See Supplementary Figure 1 for plots of BRCA1, TET2, PPM1D, LDLR, EML2 and DEDD2.

**Table 1.** Survival analysis results for genes that exceeded exome-wide significance in at least one of the six survival phenotypes analyzed in this study. ^a^Number of individuals who carry at least one PTV, ^b^Number of unique PTVs,^c^ hazard ratio, ^d^P value, ^e^see Supplementary Table 2 for full list of significant phenotypes

| Gene | Survival phenotype | N PTV carriers<sup>a</sup> | N PTVs<sup>b</sup> | HR<sup>c</sup> | P<sup>d</sup> | Select significant PheWAS phenotypes<sup>e</sup> |
| --- | --- | --- | --- | --- | --- | --- |
| BRCA2 | All proband | 675 | 191 | 2.57 | 5.02E-15 | ICD10 Z40.0 Prophylactic surgery for risk-factors related to malignant neoplasms; ICD10 Z80.3 Family history of malignant neoplasm of breast; ICD10 Z85.3 Personal history of malignant neoplasm of breast; ICD10 Z90.1 Acquired absence of breast(s); Illness of mother: Breast cancer |
|  | Females | 367 | 140 | 3.12 | 1.68E-10 |  |
|  | Males | 308 | 122 | 2.25 | 8.43E-07 |  |
|  | Combined parents | 668 | 190 | 1.60 | 2.30E-22 |  |
|  | Mothers | 658 | 186 | 1.60 | 4.70E-26 |  |
|  | Father | 652 | 186 | 1.21 | 1.35E-05 |  |
| BRCA1 | All proband | 258 | 100 | 2.59 | 2.07E-06 | ICD10 Z40.0 Prophylactic surgery for risk-factors related to malignant neoplasms; ICD10 Z90.1 Acquired absence of breast(s); ICD10 Z85.3 Personal history of malignant neoplasm of breast; ICD10 Z80.3 Family history of malignant neoplasm of breast; ICD10 Z80.4 Family history of malignant neoplasm of genital organs |
|  | Females | 124 | 61 | 4.38 | 1.01E-06 |  |
|  | Males | 134 | 66 | 1.94 | 1.35E-02 |  |
|  | Combined parents | 251 | 98 | 1.41 | 5.26E-05 |  |
|  | Mothers | 247 | 97 | 1.64 | 5.35E-11 |  |
|  | Father | 245 | 97 | 1.04 | 0.58 |  |
| TET2 | All proband | 675 | 440 | 1.75 | 8.60E-07 | ICD10 D46.9 Myelodysplastic syndrome, unspecified; ICD10 D70 Agranulocytosis; Eosinophill count; Eosinophill percentage; Platelet distribution width; Neutrophill count; White blood cell (leukocyte) count |
|  | Females | 327 | 245 | 1.38 | 0.15 |  |
|  | Males | 348 | 255 | 1.92 | 6.44E-07 |  |
|  | Combined parents | 665 | 434 | 0.95 | 0.22 |  |
|  | Mothers | 655 | 429 | 0.95 | 0.28 |  |
|  | Father | 644 | 423 | 0.97 | 0.43 |  |
| EML2 | All proband | 355 | 46 | 2.46 | 8.82E-07 | None |
|  | Females | 212 | 32 | 1.52 | 0.21 |  |
|  | Males | 143 | 29 | 3.35 | 3.33E-08 |  |
|  | Combined parents | 350 | 46 | 0.95 | 0.44 |  |
|  | Mothers | 345 | 44 | 1.03 | 0.70 |  |
|  | Father | 336 | 45 | 0.88 | 3.89E-02 |  |
| PPM1D | All proband | 135 | 59 | 3.14 | 7.70E-09 | ICD10 D47.1 Chronic myeloproliferative disease; ICD10 Z51.1 Chemotherapy session for neoplasm; Mean corpuscular haemoglobin; Self-reported medication: hydroxycarbamide; Red blood cell (erythrocyte) count; Mean corpuscular volume |
|  | Females | 60 | 34 | 4.10 | 8.48E-06 |  |
|  | Males | 75 | 45 | 2.73 | 8.07E-05 |  |
|  | Combined parents | 133 | 59 | 1.04 | 0.69 |  |
|  | Mothers | 131 | 59 | 1.00 | 0.97 |  |
|  | Father | 129 | 58 | 1.08 | 0.42 |  |
| LDLR | All proband | 51 | 28 | 1.51 | 0.48 | ICD10 E78.0 Pure hypercholesterolaemia; ICD10 I25.1 Atherosclerotic heart disease; Self-reported illness: high cholesterol Self-reported medication: ezetimibe; Self-reported medication: rosuvastatin; Self-reported medication: atorvastatin; Self-reported medication: aspirin; Self-reported medication: lipitor 10mg tablet |
|  | Females | 33 | 20 | 0.81 | 0.84 |  |
|  | Males | 18 | 12 | 2.68 | 0.16 |  |
|  | Combined parents | 51 | 28 | 2.28 | 5.14E-07 |  |
|  | Mothers | 51 | 28 | 1.62 | 1.97E-03 |  |
|  | Father | 50 | 28 | 1.60 | 2.11E-03 |  |
| DEDD2 | All proband | 15 | 9 | 4.75 | 0.028 | None |
|  | Females | 6 | 5 | 5.16 | 0.10 |  |
|  | Males | 9 | 6 | 4.35 | 0.14 |  |
|  | Combined parents | 15 | 9 | 2.66 | 1.19E-03 |  |
|  | Mothers | 15 | 9 | 4.20 | 2.28E-07 |  |
|  | Father | 15 | 9 | 1.19 | 0.55 |  |

For *BRCA2, BRCA1* and *LDLR*, the majority of PTVs identified in this study had been described earlier as causing monogenic disease. Specifically, 139 out of 190 *BRCA2* PTVs, and 73 out of 100 *BRCA1* PTVs were reported as Pathogenic or Likely pathogenic in Clinvar (version 20200316)^17^ for hereditary breast and ovarian cancer syndrome (Supplementary Table 1). Similarly, 24 out of 28 *LDLR* PTVs are reported as Pathogenic or Likely pathogenic for familial hypercholesterolemia. Of the remaining four genes, only PTVs in *PPM1D* have been linked earlier to disease, with two out of the 59 *PPM1D* PTVs identified in this study reported to cause a neurodevelopmental syndrome (Supplementary Table 1).

To gain insights into the disease endpoints and biological processes that drive associations with lifespan, we performed PTV-burden phenome-wide association studies (PheWAS) for each of the seven exome-wide significant genes across 4,130 semi-automatically derived UK Biobank phenotypes. Consistent with their established roles in hereditary breast, ovarian and prostate cancers^18^, *BRCA2* and *BRCA1* PTV-burden were associated with increased risk of breast (*BRCA2*: odds ratio, OR=4.86, P=2.41e-37; *BRCA1*: OR=10.11, P=1.53e-34) and ovarian cancer (*BRCA2*: OR=10.31, P=1.11e-16; *BRCA1*: OR=14.77, P=1.14e-8) in females, while *BRCA2* was associated with prostate cancer in males (OR=3.36, P=8.20e-10) (Supplementary Table 2, Supplementary Figure 2). *TET2* mutations are a known cause of hematopoietic abnormalities^19^. Consistently, PheWAS revealed *TET2* PTV-burden as associated with reduced eosinophil (β=-0.42 SDs, P=2.44e-30) and neutrophil count (β=-0.29 SDs, P=9.13e-14), along with an increased risk for myelodysplastic syndrome (OR=16.06, P=9.59e-37) and agranulocytosis (OR=4.29, P=4.78e-13). We also observed male-specific associations between *TET2* PTV-burden with thrombocytopenia (OR=7.15, P=4.63e-11 in males versus P=0.46 in females) and the risk for acute myeloid leukemia risk (OR=12.01, P=2.39e-9 in males versus P=0.20 in females), a sex bias that matches findings from observational studies which showed both lower mean platelet counts and a higher prevalence of these diseases in men than in women^20,21^. The *PPM1D* PTV-burden PheWAS identified associations with increased mean corpuscular hemoglobin (β=0.81, P=7.91e-9), chronic myeloproliferative disease (OR=29.35, P=2.22e-6) and hydroxycarbamide medication usage (OR=41.53, P=4.49e-7), reflecting previous observations that PPM1D-truncating mutations are enriched in myeloproliferative neoplasms and confer resistance to chemotherapy^22,23^. The phenotypes associated most strongly with *LDLR* PTV-burden include self-reported high cholesterol (OR=19.79, P=1.53e-19), pure hypercholesterolemia (OR=7.66, P=2.85e-10), chronic ischemic heart disease (OR=7.59, P=3.52e-6) and rosuvastatin medication usage (OR=31.97, P=1.93e-11), consistent with results from previous rare variant-burden studies of *LDLR* for cardiovascular phenotypes^24^. PTV-burden PheWAS did not reveal any significantly associated phenotypes for *DEDD2* or *EML2. DEDD2* encodes death effector domain-containing protein 2 with roles in apoptosis and tumor growth^25, 26^, while the molecular functions and disease relevance of *EML2* is unclear.

In addition to a gene-based approach for exploring the genetic basis of human lifespan, we next combined PTVs across genes annotated for similar functions and conducted gene-set burden survival analyses. Using gene-set definitions from 4,589 ConsensusPathDB pathways^27^, we identified 29 pathways that were significantly associated with survival at a 5% Bonferroni threshold (P<1.09e-5). All significant pathways were linked to cancer susceptibility and included *homologous recombination* (Wikipathways; HR=1.80, P=8.38e-14), *role of brca1, brca2 and atr in cancer susceptibility* (BioCarta; HR=1.42, P=2.28e-12) and *p53 pathways* (Protein Interaction Database; HR=1.24, P=4.80e-7), among others (Supplementary Table 3, Supplementary Figure 3). Twenty pathways remained nominally significant (P < 0.05) after excluding the seven genes from PTV-burden survival analyses along with *ATM*, a member of 27 of the 29 pathways and in single-gene analyses slightly below the exome-wide significance cutoff (HR=1.89, P=5.44e-6). This suggests that additional members of these pathways will reach gene-level significance when sample sizes for PTV-burden analyses increase further.

The overlap between our PTV-burden with previous GWAS of ageing phenotypes results was modest. Of the seven genes associated with survival in our PTV-burden analyses, only variants near *LDLR* has also been identified through GWAS. Conversely, of the 22 other reported GWAS loci^1^ only *MICB* (HR=2.23, P=4.63e-5) and *SEMA6B* (HR=1.81, P=4.56e-3) were nominally significant in at least one of the six survival phenotypes analyzed in this study (Supplementary Table 4).

Our results reflect the demographic makeup of the UK Biobank and may not fully extrapolate to other populations. The median age at the censoring date in our WES sample was 67, while the median age at death in the UK is 82 and 85 years for males and females respectively^28^. As such, causes of death that do not typically affect middle-aged individuals are likely underrepresented. Moreover, UK Biobank participants are healthier than the general UK population, with participants being less likely to smoke, be obese or drink^29^, potentially diluting our ability to capture the effects of these factors on mortality. Using parental lifespan as a proxy phenotype reduces these ascertainment biases, although compared to directly observed phenotypes this approach requires much larger sample sizes for reaching similar statistical power^30^. Genetic associations with parental lifespan that were not detected in probands may also reflect recent advances in medical care and other environmental factors. For instance, the widespread use of statin medications since the 1990s to reduce the risk of heart disease^31^ may partly explain why *LDLR* PTV-burden association with survival was only observed in the parents of UK Biobank participants.

UK Biobank demographics cannot fully account for the limited overlap between our PTV-burden results and loci identified by previous ageing GWAS, which also included UK Biobank participants^7–9^. Instead, GWAS signals might be driven by mechanisms unrelated to protein loss of function, or act via other genes at the same locus or *in trans*. Also, at current sample sizes PTV-based burden analyses remain underpowered to detect associations for many genes due to a lack of observations in populations that are deprived from high-impact loss-of-function variants due to purifying selection. Nevertheless, our analyses identified robust signals for genes to impact lifespan that GWAS were yet unable to detect, such as for *BRCA2*, demonstrating the potential for gene-based analyses in large sequenced cohorts to complement common variant GWAS.

In conclusion, using WES data from 238,239 UK Biobank participants, we identified seven genes for which loss-of-function is associated with human lifespan. Future efforts may expand our approach to gain-of-function and other rare protein-coding alleles and incorporate additional factors that impact a person’s age at death which in most cases will be a reflection of their past health and lifestyle. Our study establishes the importance of individual genes and pathways on human lifespan at the population level and highlights intervention points that, if adequately addressed, may allow for greater wellbeing as we age.

## Online Methods

### UK Biobank

The UK Biobank is a prospective study of over 500,000 participants aged 40-69 years recruited from 2006-2010 in the UK^32^. Participant data include health records, medication history and self-report survey information along with imputed genome-wide genotypes^33^.

Whole-exome sequencing (WES) data for UK Biobank participants were generated at the Regeneron Genetics Center (RGC) as part of a collaboration between AbbVie, Alnylam Pharmaceuticals, AstraZeneca, Biogen, Pfizer, Regeneron and Takeda with the UK Biobank^13^. WES data were processed using the RGC SBP pipeline described in Van Hout et al., 2019^16^. RGC generated a QC-passing “Goldilocks” set of 23,482,637 genetic variants from a total of 302,331 sequenced UK Biobank participants for analysis.

### Survival phenotypes

UK Biobank participants’ age at death were automatically linked through NHS Digital (for England and Wales) and Information and Statistics Division (for Scotland)^34^. English/Welsh and Scottish death records were current as of January 31, 2018 and November 30, 2016, respectively. We used these dates as the censoring dates for participants depending on whether the individual listed their home area in either England/Wales or Scotland. For individuals without a death record, we assumed they were alive on the censoring date, and calculated their current age to be January 2018 minus their month and year of birth for those residing in England/Wales, or November 2016 minus their month and year of birth for those residing in Scotland.

During the initial UK Biobank assessment between 2006 to 2010, all participants were asked to record the ages of their father/mother if alive, or their age at death. Participants were also asked whether they were adopted as a child. Repeat assessment (2012 onwards) data were available for 56,378 participants. We extracted the parental ages/ages at death from the most recently provided assessment visit of each participant.

### PTV annotation

Variants identified through WES were annotated with VEP v96^14^ and the LOFTEE^12^ plugin. LOFTEE applies a range of filters on stop-gained, splice-site disrupting and frameshift variants in order to exclude putative PTVs due to variant annotation and sequencing mapping errors that are unlikely to significantly disrupt gene function. For instance, stop-gained and frameshift variants that are within 50kb of the end of the transcript will be flagged as “low-confidence”. We extracted variants predicted as PTVs, flagged as “high confidence” by LOFTEE and with minor allele frequency <1% for each canonical transcript (as defined in Ensembl). In total, 572,780 unique PTVs across 19,094 genes were available for analysis. To account for relatedness, we excluded from analyses one member (at random) from each pair of ≤2^nd^ degree relatives, as well as non-white British ancestry individuals based on principal components analysis of the public genotype data^33^.

### Survival analysis

A total of 238,239 individuals were available for analysis, of which 9,405 were deceased with their age at death recorded at the censoring dates. For the parental survival analysis, we adopted a similar approach to previous studies using the UK Biobank parental information^7^ and further excluded adopted individuals (n=3,090), individuals who did not know/did not answer the parental age/age at death questions (7,383 fathers, 3517 mothers), or when a parent had died before the age of 40 (4,174 fathers, 2,571 mothers). The survival analysis in fathers included 235,149 individuals and 178,443 events, the survival analysis in mothers included 234,967 individuals and 145,281 events. We created a combined father and mother age by summing the reported ages of the parents and defining events as whether both parents were deceased (n=225,701, n events =128,045).

For each gene, we applied a Cox proportional hazard model (right censored) to test for an association between survival and PTV-burden: *h*(*x_i_*) = *h*_0_(*x_i_*)exp (*β_g_i__*, + *γ_i_**Z_i_***), where for each individual *i, x* is the age, *h*_0_ is the baseline hazard, *g* is the genotype with coefficient *β, **Z*** is a matrix of covariates with coefficients *γ*. For each gene, *g_i_* = 1 if individual *i* carries at least one PTV, otherwise, *g_i_*=0. We included baseline age, sex and 10 PCs as covariates in all the survival phenotypes analyzed, except for the proband sex-specific analyses, where the sex covariate was dropped. All Cox regressions were performed in R with the “survival” package^35^. Six survival phenotypes were analyzed: proband age, male proband age, female proband age, combined parental age, father’s age and mother’s age. Approximate exome-wide significance was defined as P<0.05/20,000 genes = 2.5e-6. As the Cox model may produce biased type I error estimates when the number of observed events/predictors are low^36^, we excluded genes with fewer than 10 PTVs among PTV carriers (for comparison purposes, exceptions to this rule were applied to *LDLR* and *DEDD2* in the proband analyses since those genes showed significant association with parental survival).

### PheWAS analysis

For genes that exceeded exome-wide significance in the Cox analysis, we performed a PTV-burden PheWAS across 4,130 semi-automatically derived UK Biobank phenotypes. Binary phenotypes included ICD10 codes (each primary ICD10, secondary ICD10 and cause of death ICD10 code was combined into a single ICD10 phenotype), self-reported health outcomes, medication usage, surgery/operation codes, and family history (fathers’ illnesses, mothers’ illnesses and siblings’ illnesses were combined into a single phenotype for each of the 12 family history illnesses ascertained for in UK Biobank questionnaires). Additional disease endpoints (e.g. breast cancer, ovarian cancer, type 2 diabetes) were derived manually by combining the ICD codes, self-report, medication, operation codes and other relevant UK Biobank fields. Quantitative phenotypes include 31 blood count phenotypes (e.g. lymphocyte count), 30 blood biochemistry phenotypes (e.g. cholesterol), 47 infectious disease antigen assays (e.g. L1 antigen for HPV), and over 400 physical (e.g. hand grip strength) and cognitive (e.g. numeric memory) measurements.

We excluded binary phenotypes with fewer than 100 cases among the 238,239 post-QC set of UK Biobank individuals. PTV burden testing for binary phenotypes were performed using logistic regression in all individuals as well as males and females separately. For gene-phenotype pairs where the PTV burden P-value < 0.05, we repeated the analysis using the Firth method to account for situations where the logistic regression outputs may be biased due to separation^37^. For quantitative phenotypes, we excluded phenotypes with fewer than 500 observations. For each phenotype, outlying individuals (defined as having > 5 SDs from the mean) were excluded. Burden testing was performed using linear regression on both the raw and inverse rank normal transformed quantitative phenotypes in all individuals as well as males and females separately. In both the logistic and linear models, we included covariates age, sex and 10 PCs. Sex was excluded as a covariate for the sex-specific analyses. We defined a 5% Bonferroni-corrected phenome-wide significance threshold of P<1.2e-5 (=0.05/4,130).

### Gene-set PTV survival analysis

We performed gene set PTV survival analysis by first grouping genes into pathways as defined by ConsensusPathDB-human release 30^27^, which integrates over 4,589 pathways from 32 databases including KEGG, BioCarta and WikiPathways. For each pathway, we collapsed PTVs from all genes that are annotated as being members of a respective pathway into a single score and tested this score for association with survival using the same Cox model approach as described above for individual genes.

## Acknowledgements

This research has been conducted using the UK Biobank Resource under Application Numbers 25570 and 26041. We thank all the participants and researchers of UK Biobank for making these data open and accessible to the research community. The Regeneron Genetics Center and consortium members AbbVie, Alnylam Pharmaceuticals, AstraZeneca, Biogen, Pfizer, Regeneron and Takeda are acknowledged for generation and initial quality control of the whole-exome sequencing data. We thank Eric Marshall, Yongsheng Huang and Frank Nothaft for infrastructure support.

## Notes

### Competing Interest Statement

Jimmy Z. Liu, Chia-Yen Chen, Ellen A. Tsai, Christopher D. Whelan, David Sexton, Sally John and Heiko Runz are employees of Biogen

