## Supplementary figures for "The burden of rare protein-truncating genetic variants on human lifespan"

Supplementary Figure 1. Kaplan-Meier plots for six survival phenotypes comparing survival rates of PTV carriers and non-carriers at *BRCA1*, *TET2*, *EML2*, *PPM1D*, *LDLR* and *DEDD2.* Each cross represents a right censored observation. The shaded areas represent the 95% confidence interval of the curve.

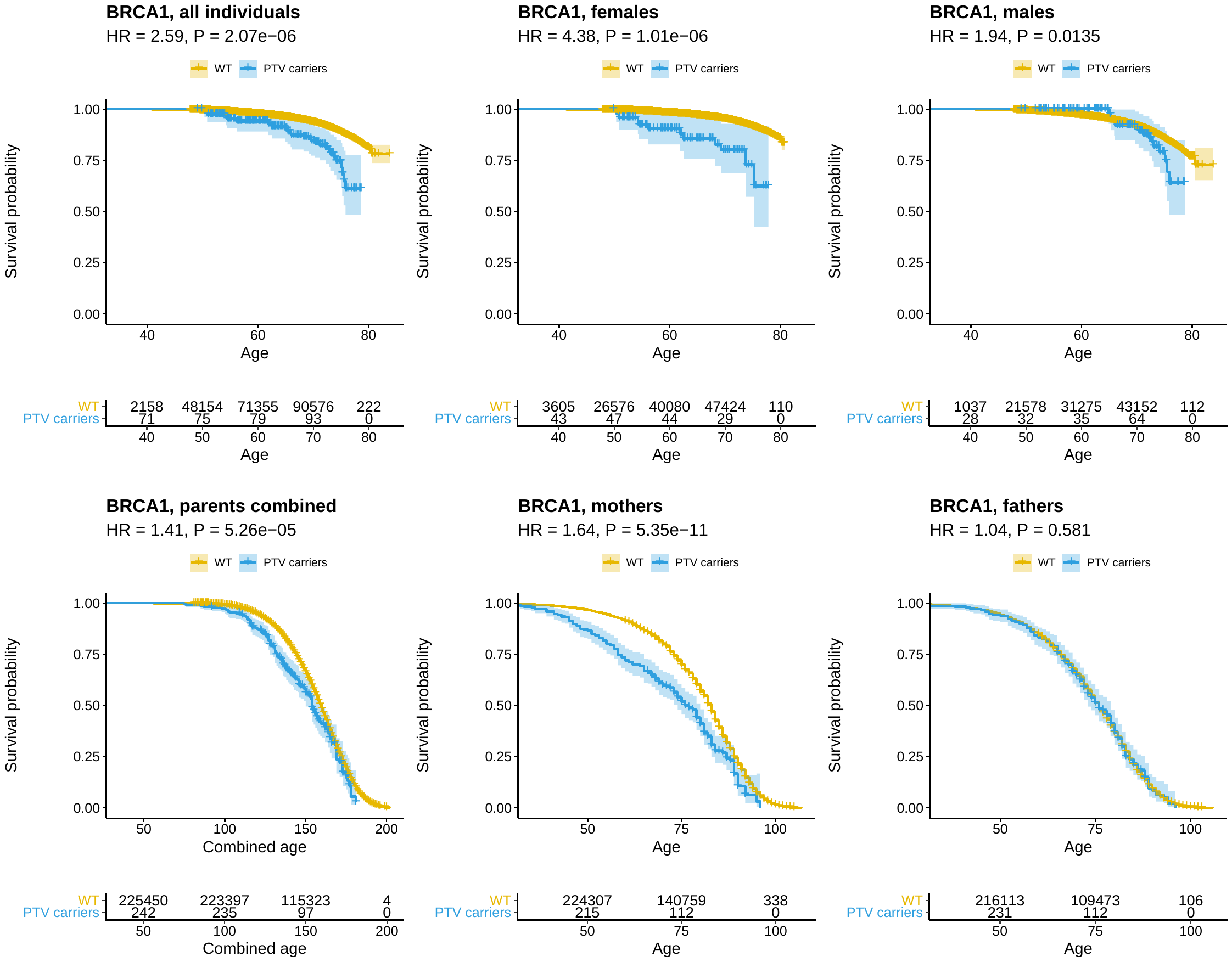

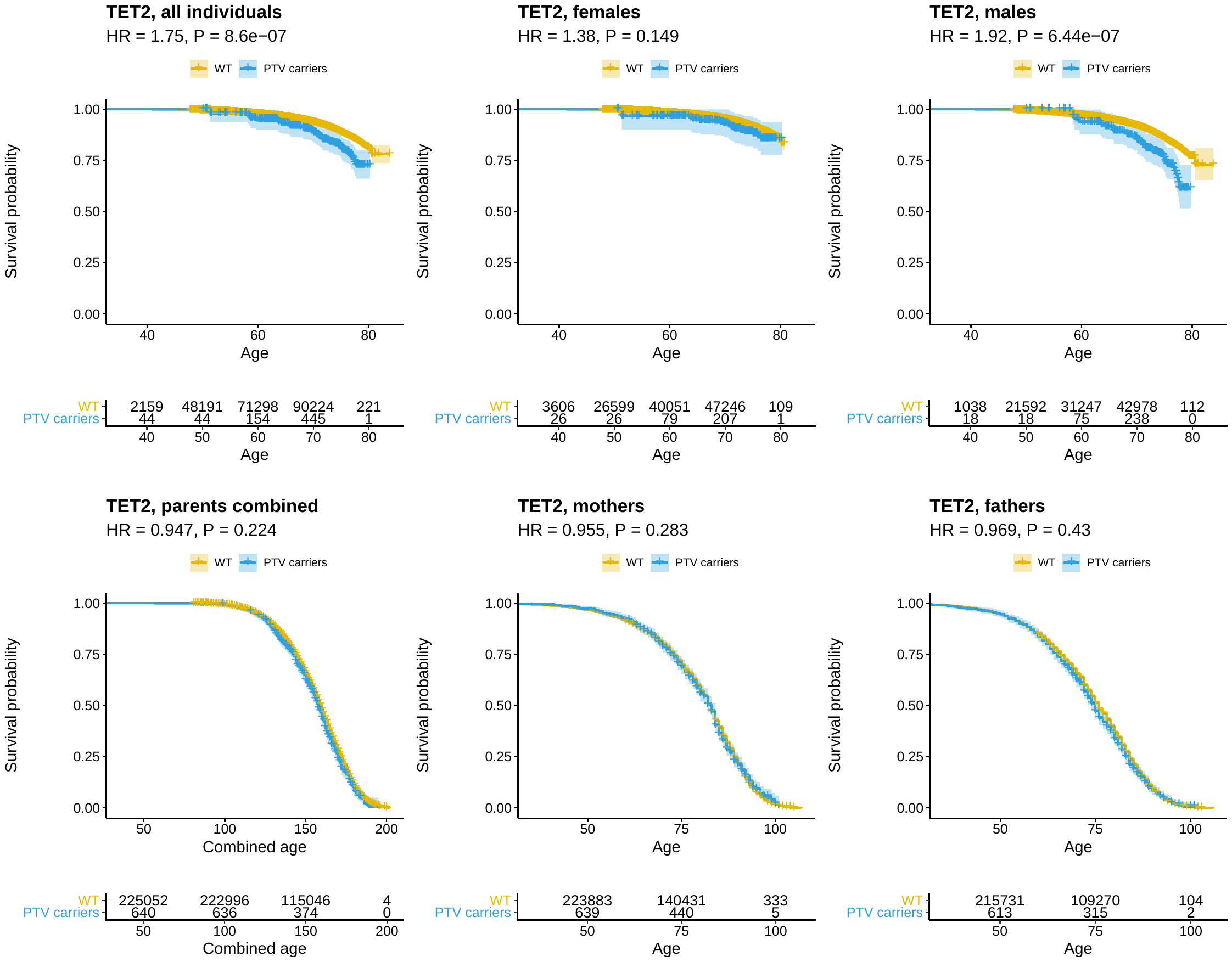

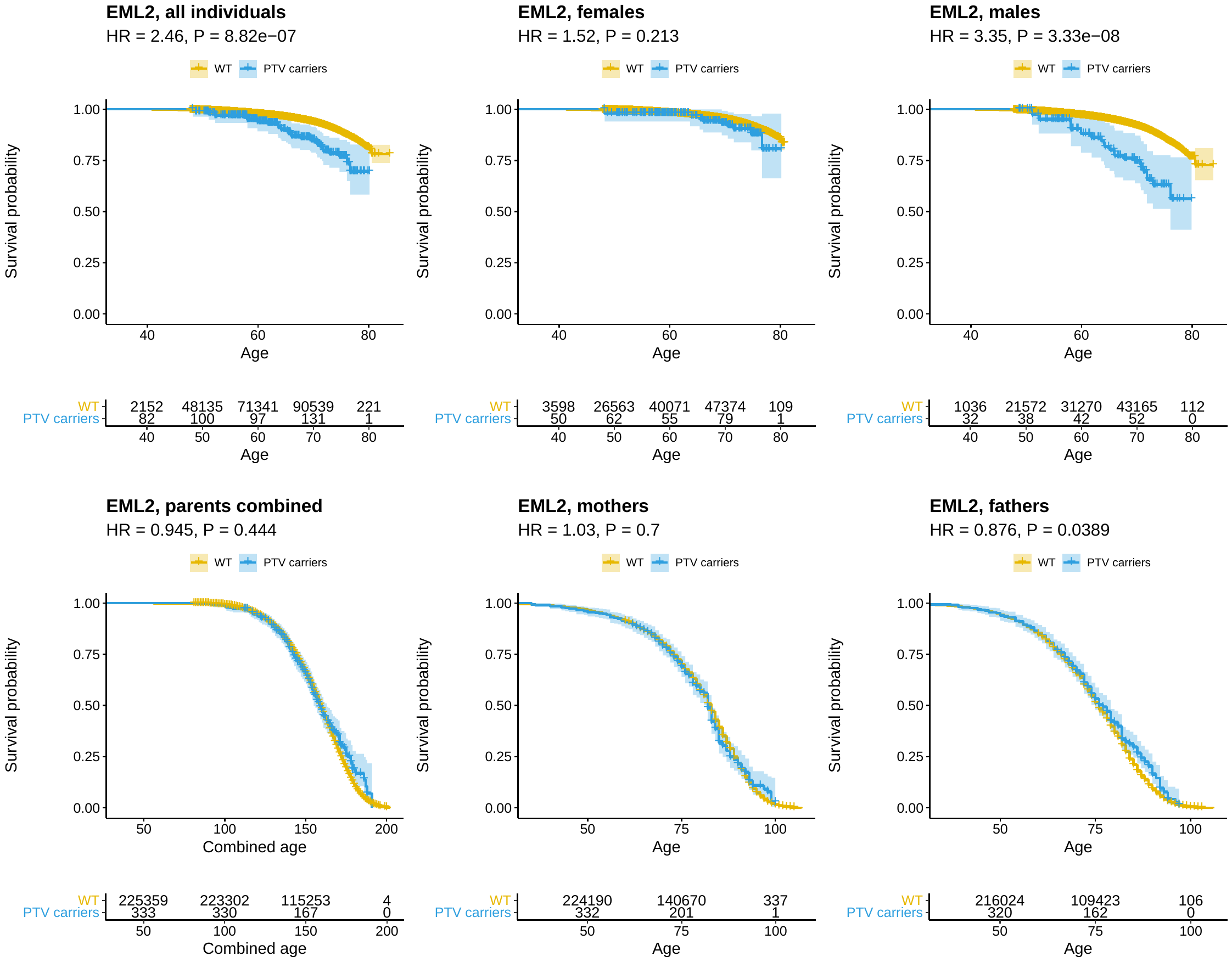

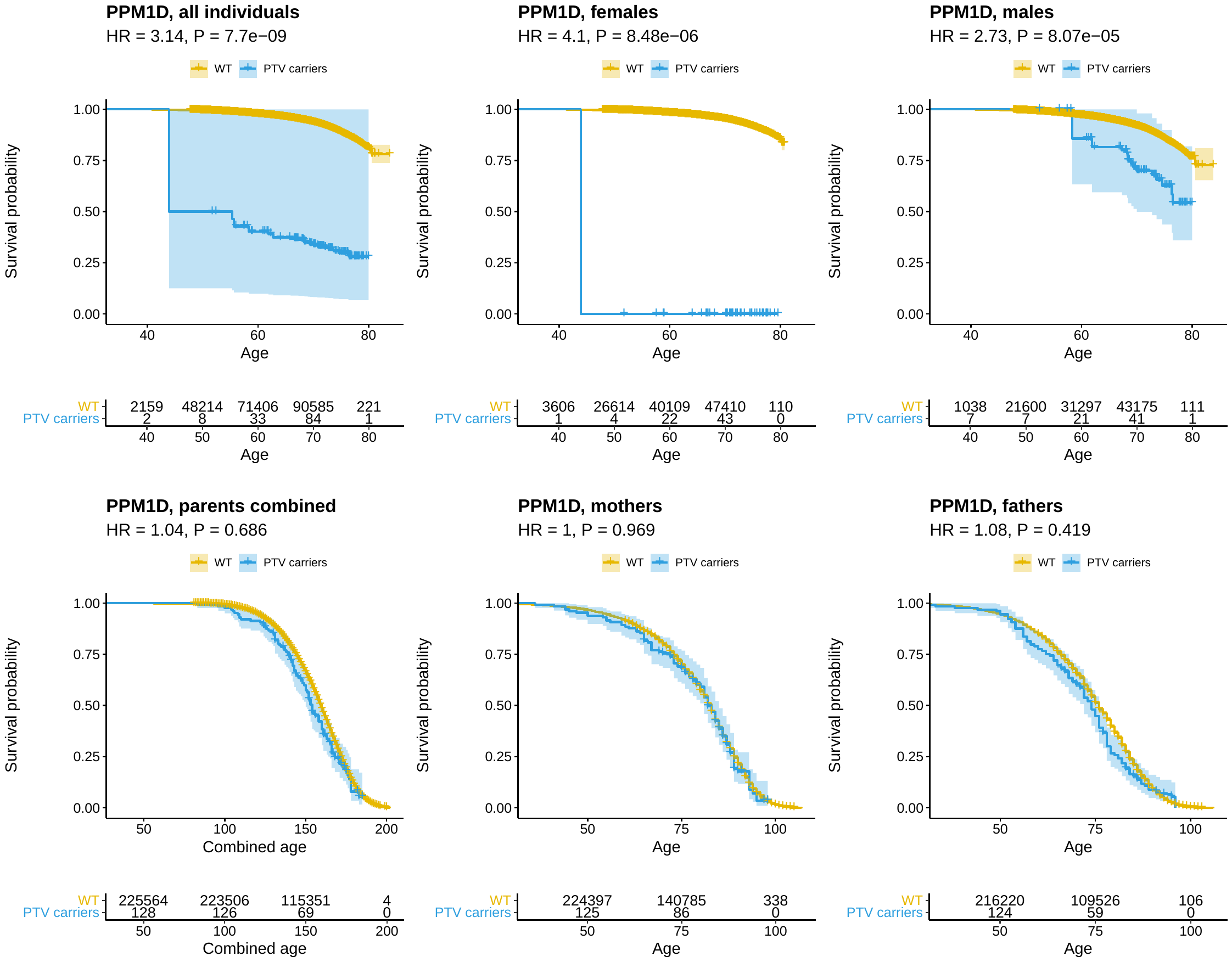

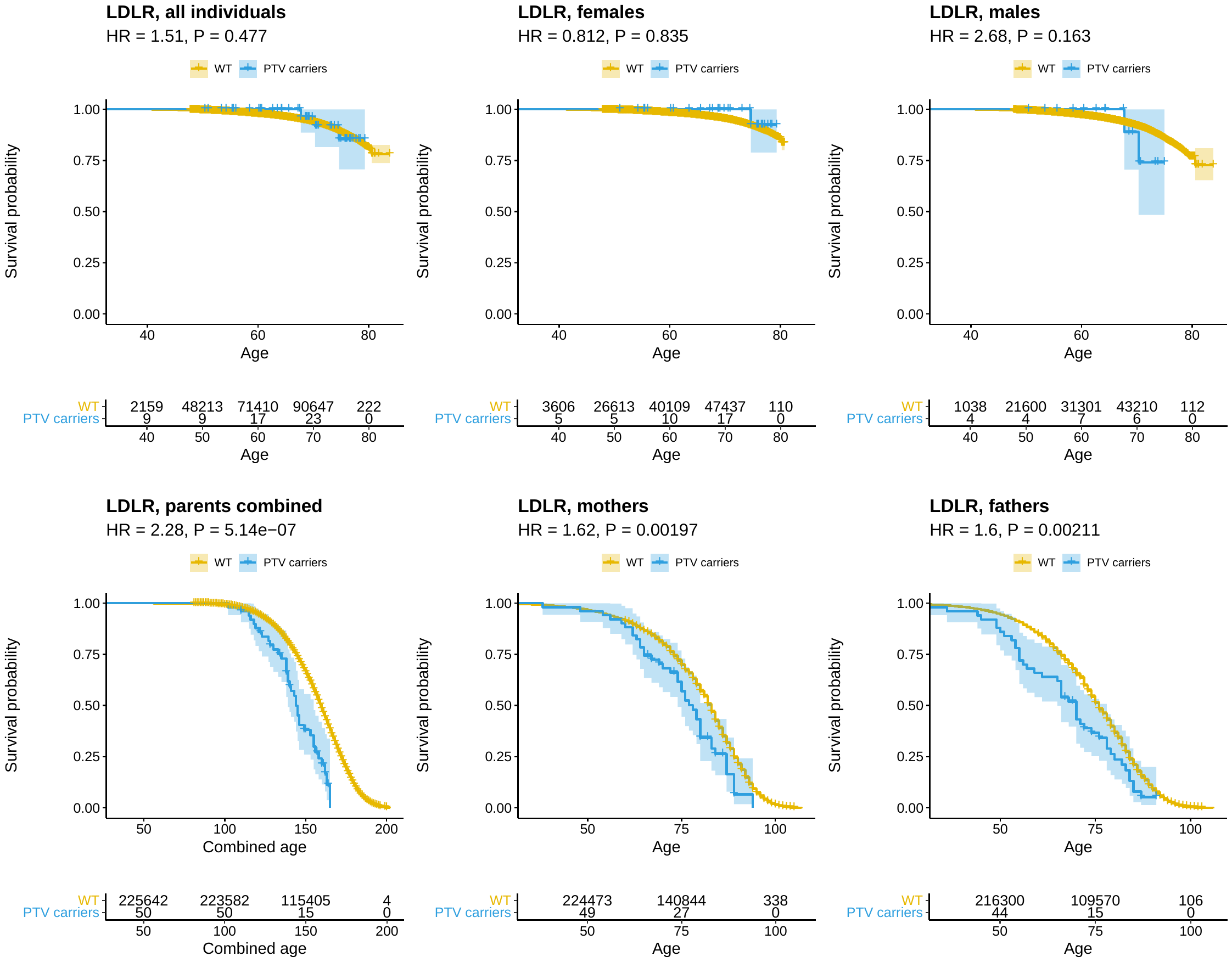

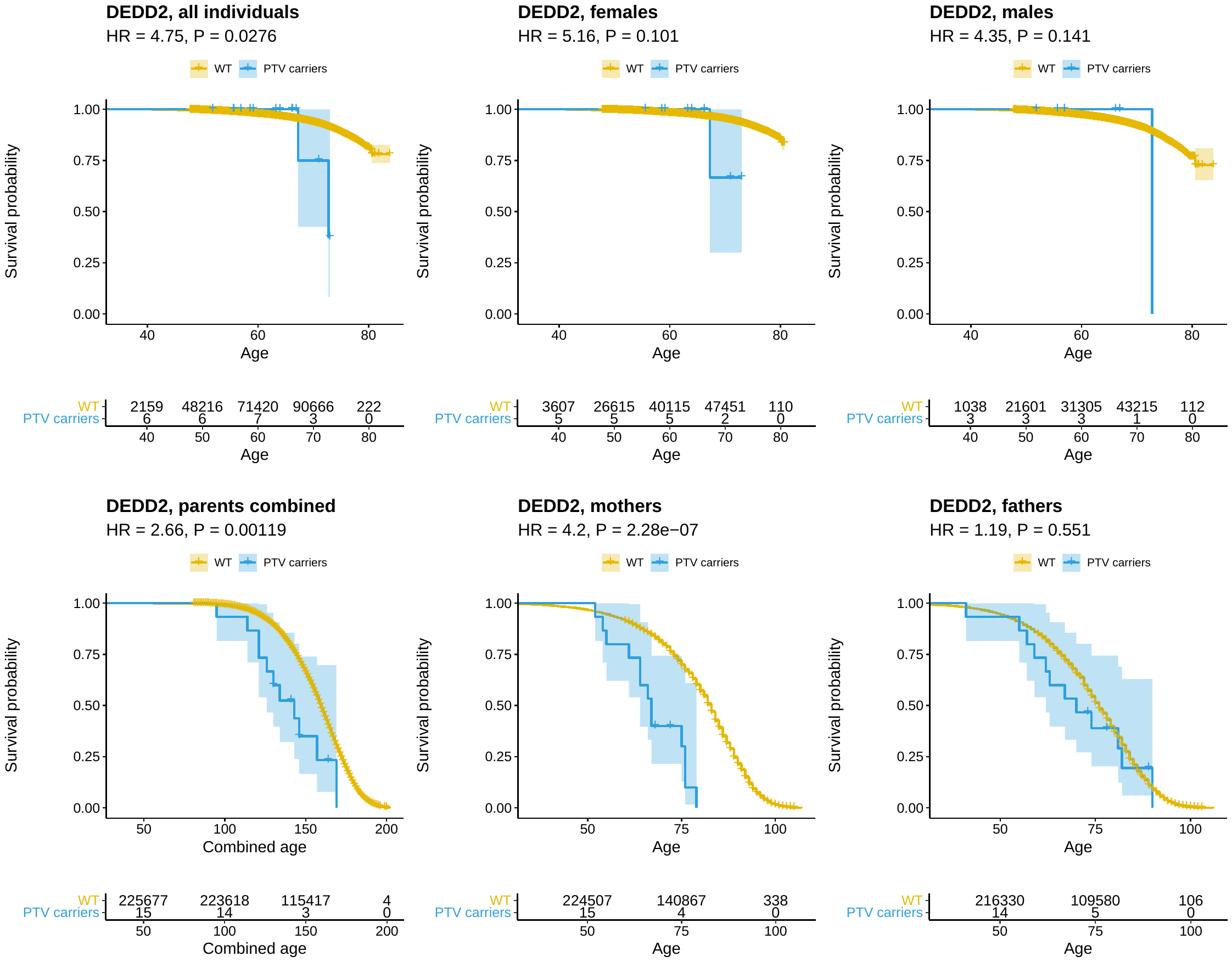

Supplementary Figure 2. Manhattan plots of PheWAS results for *BRCA2*, *BRCA1*, *TET2*, *PPM1D*, *LDLR*, *EML2* and *DEDD2* across 4,130 phenotypes. Each point represents a phenotype. The y-axis denotes the strength of association between the phenotype and gene PTV-burden. Phenotype categories are colored on the x-axis. Select significant phenotypes are labeled. See Supplementary Table 2 for detailed association results.

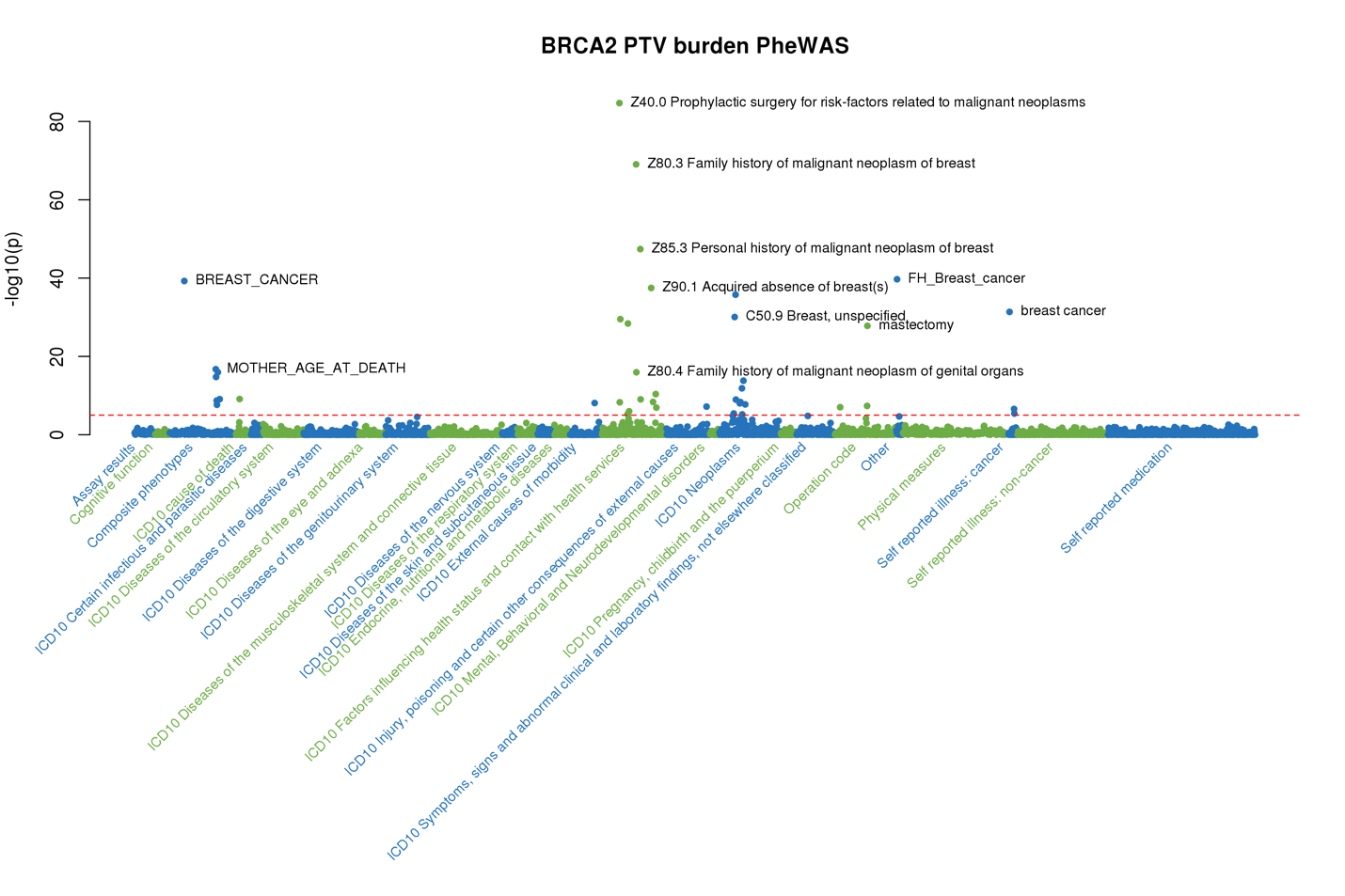

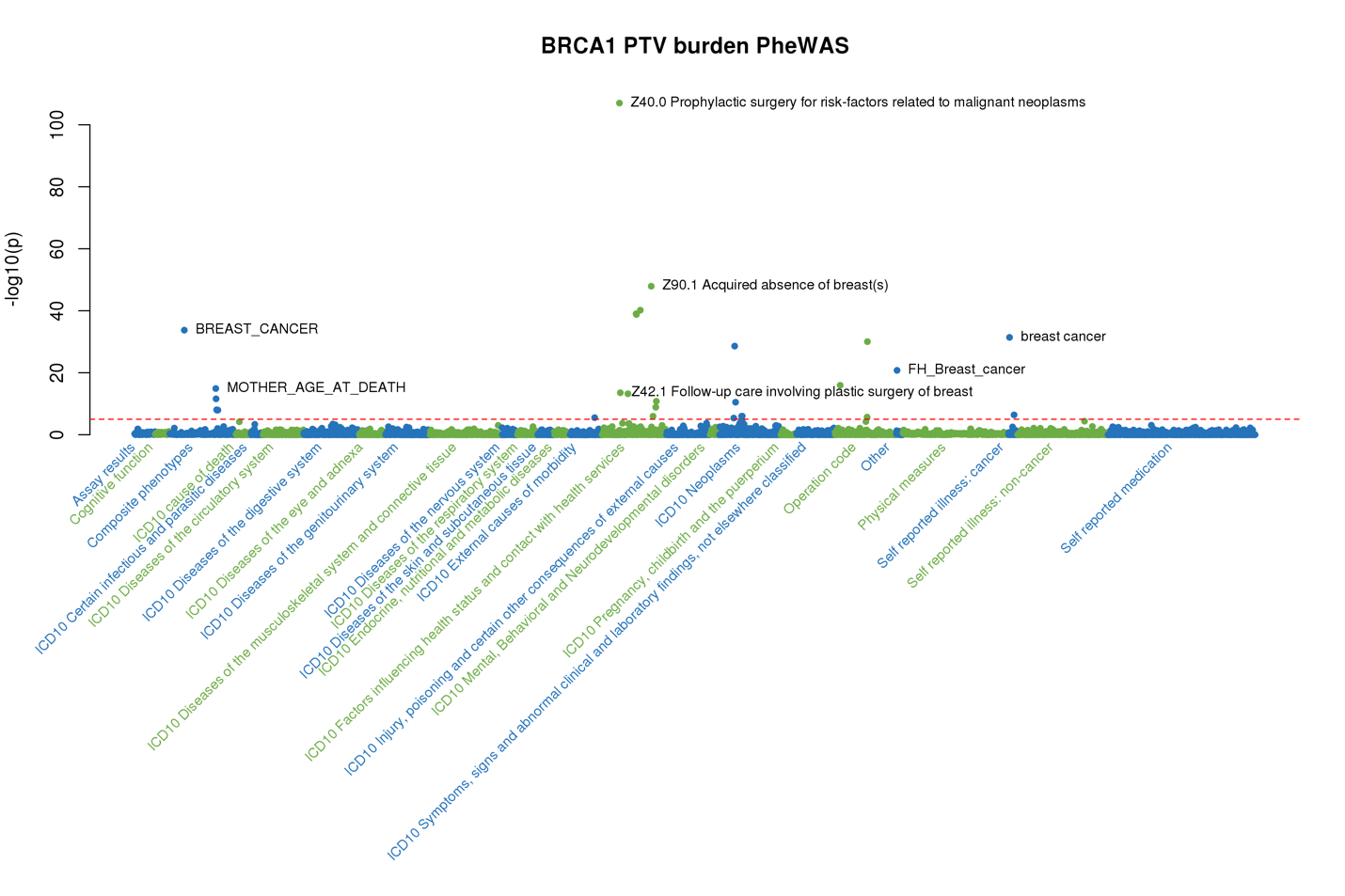

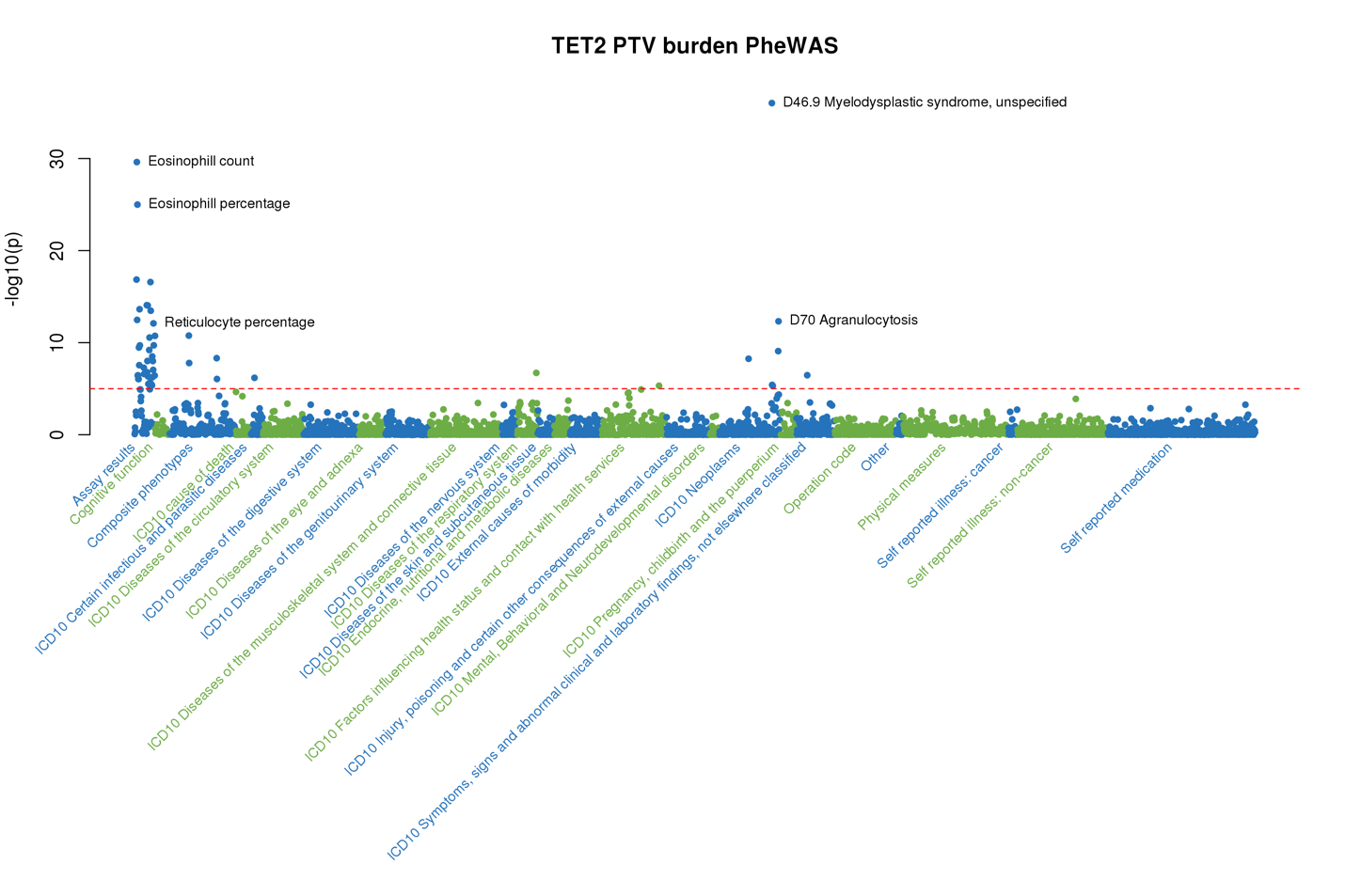

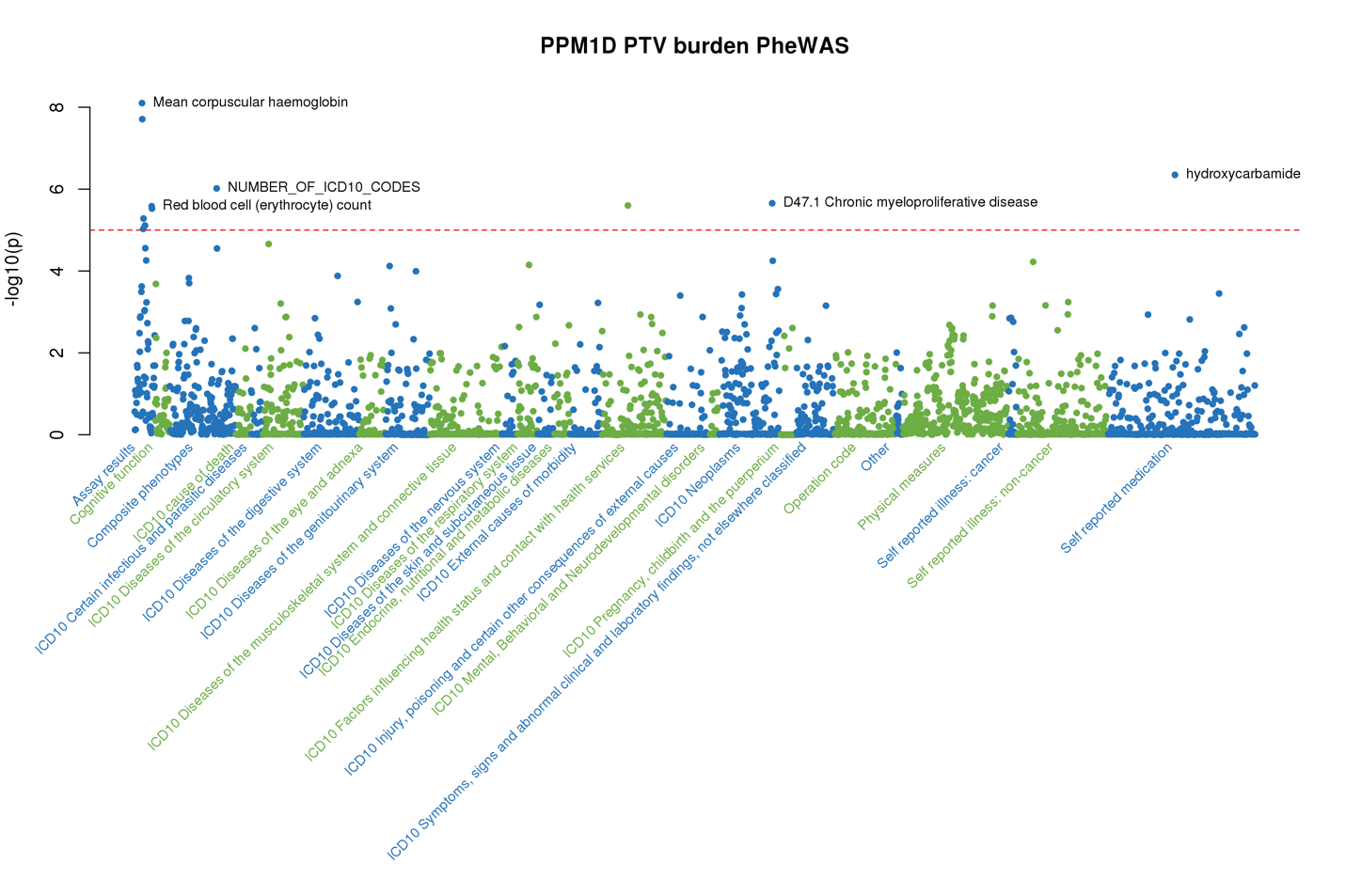

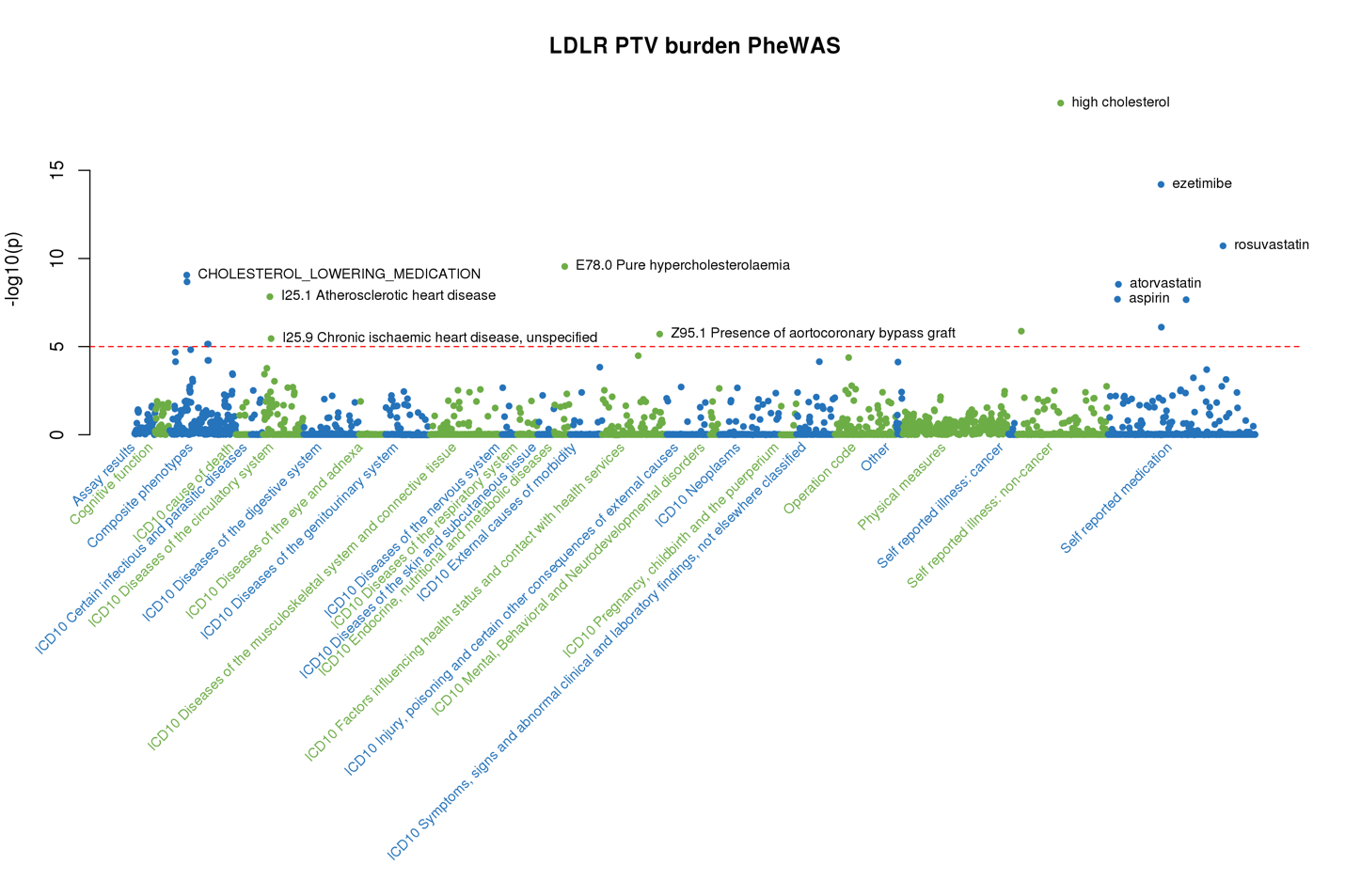

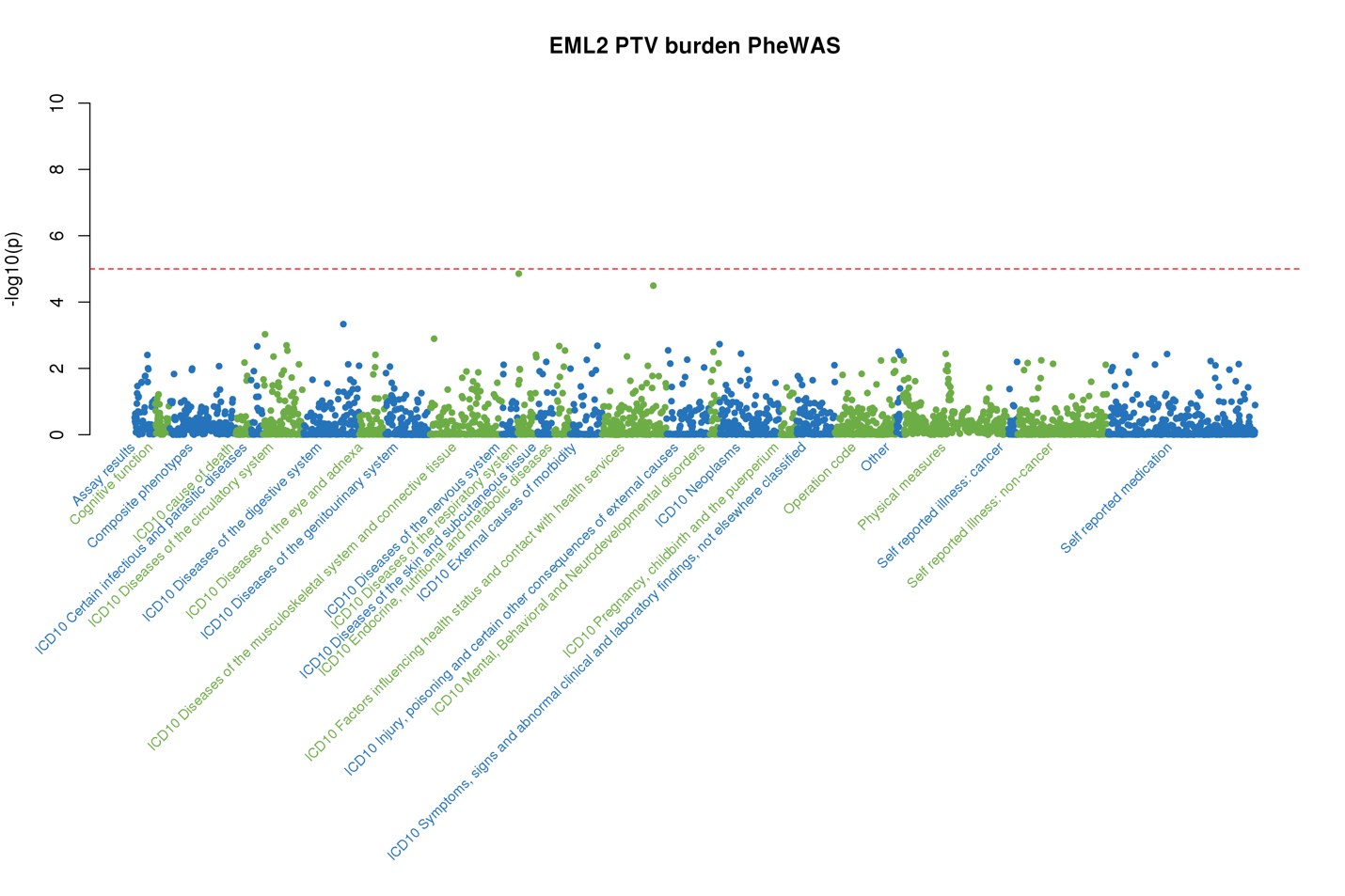

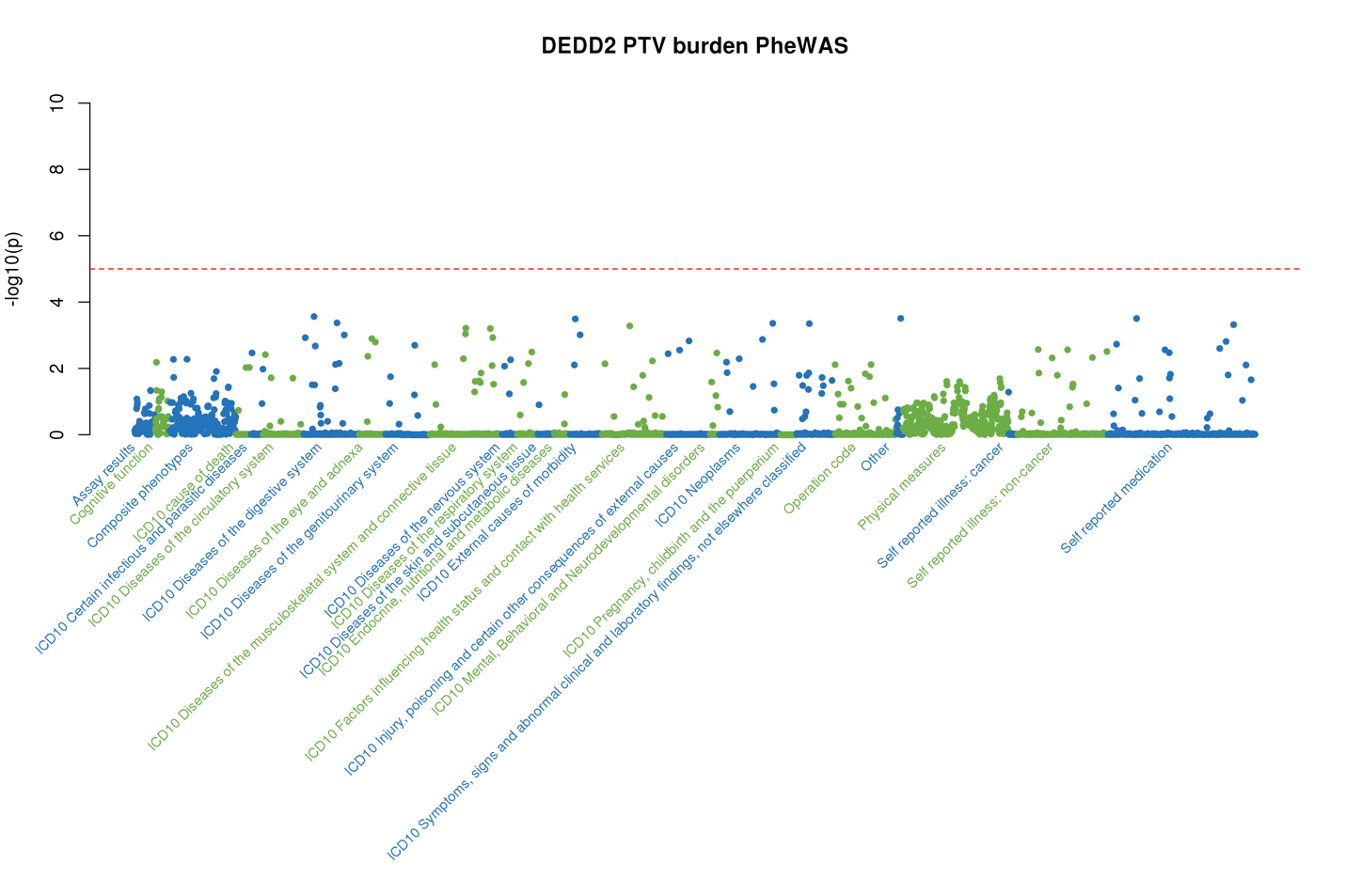

Supplementary Figure 3. Manhattan plot of gene-set PTV burden analysis results. Each point represents a gene-set. Categories are colored on the x-axis according to the source of the gene-set. Select significant gene-sets are labeled. The purple triangles in the second panel indicates pathways that were significant in first panel. See Supplementary Table 2 for detailed results.

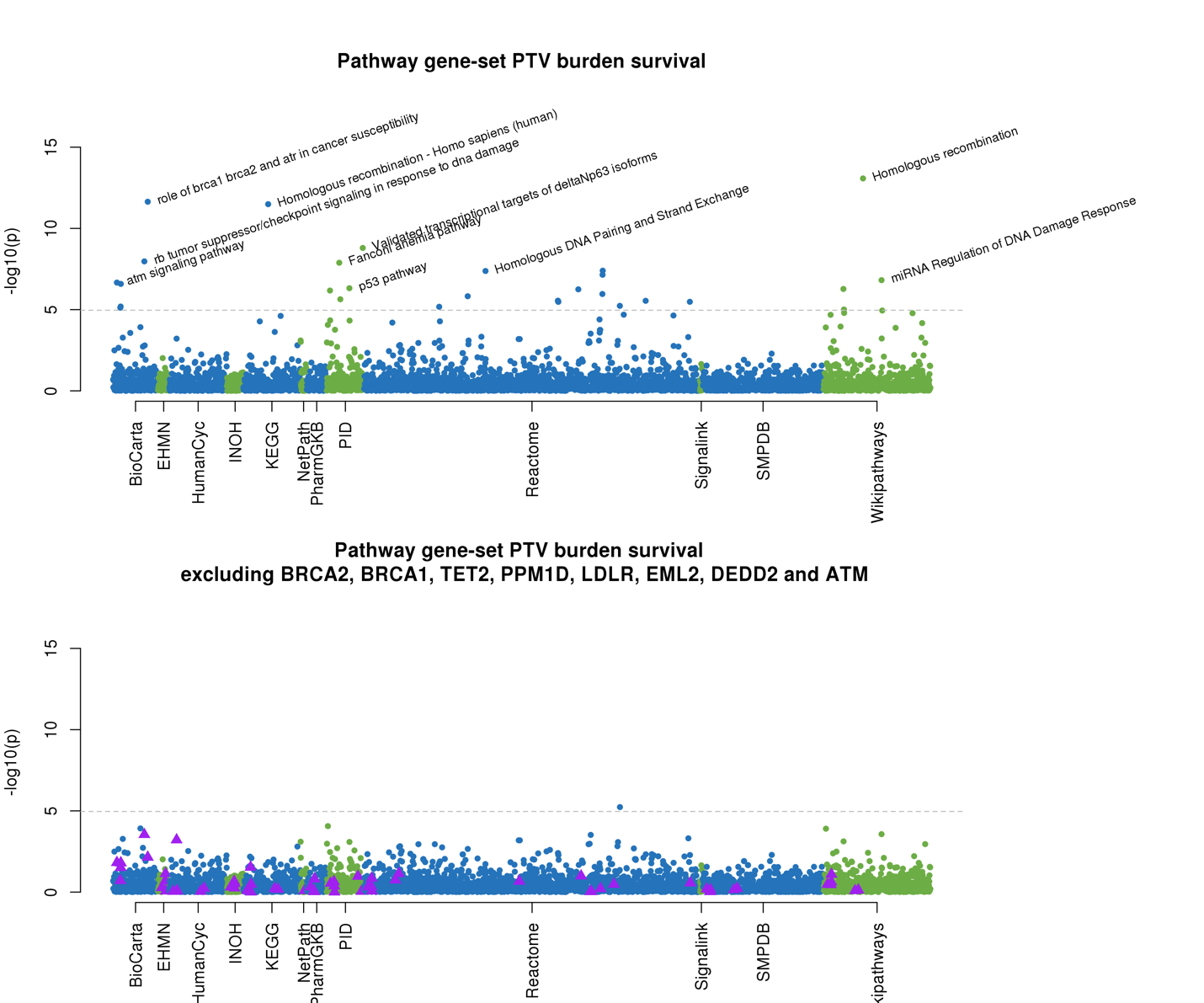
